# Effects of low intensity pulsed ultrasound stimulation on metabolic lipolysis of adipocytes

**DOI:** 10.1101/2022.03.29.486238

**Authors:** Sangnam Kim, Sangpil Yoon

## Abstract

Obesity is closely related to several metabolic diseases along with abnormal increase in fat cells. Reducing size and number of fat cells, a procedure known as lipolysis, may be used to prevent obesity as a potential therapy, which also requires fundamental understanding of the mechanisms of lipolysis at molecular level upon different types of stimulations. Here, we used low intensity pulsed ultrasound (LIPUS) stimulation to investigate underlying mechanisms of the activation of lipolysis and autophagy related genes and signaling pathways of adipocytes differentiated from 3T3-L1 cells. LIPUS with the center frequency of 2 MHz was applied for 10 minutes per day for three days. After LIPUS stimulation, quantitative reverse transcription polymerase chain reaction (RT-qPCR) and Western blot were used to determine the regulation of lipolytic factors such as adipose triglyceride lipase (ATGL), hormone-sensitive lipase (HSL), and monoacylglycerol lipase (MGL). At RNA level, all three factors were upregulated while only MGL was upregulated at protein level, which presents slightly different activation pattern of lipolysis compared to widely used chemical stimulation. These results demonstrate that LIPUS stimulation can promote the lipolytic capacity of adipocytes in the differentiated state. The differences between transcriptional genes and metabolites were analyzed by transcript analysis and metabolomic profiling experiments. Cellular RNA-sequencing (RNA-Seq) showed an increase in lipolysis and immune-related genes and autophagy related genes after LIPUS stimulation. This study may provide an important experimental basis for the clinical applications and a fundamental understanding of the mechanisms of lipolysis using LIPUS stimulation.

## 1. Introduction

Obesity is the leading cause of metabolic disease in the United States [1]. Being overweight and obese is defined as an abnormal or excessive accumulation of fat that poses a health risk. A body mass index (BMI) of 25 or more is overweight, and a BMI of 30 or more is obese. The problem has grown to an epidemic level with over 4 million deaths annually due to being overweight or obese in 2017 [2]. It was reported that overweight and obesity contribute to increase the risk of developing malignant tumors and cancer-related death in the United States [3]. At cellular and molecular level, increased size and number of fat cells in obese individuals are observed due to excess energy storage in fat cells. Obesity surgery, chemical therapy, and medical therapy are the traditional methods to treat obesity. Because of undesirable side effects of existing treatments, it is important to develop new strategies and approaches that can complement existing treatments. Therefore, efforts are needed to reduce the size and number of fat cells, also known as promoting lipolysis [4].

Lipolysis is the process by which fat is broken down in our body through enzymes and water or hydrolysis [5]. Triglycerides are glycerol derivatives that are stored as lipid droplets within the fat compartment, where lipolysis occurs. Lipolytic lipase sequentially hydrolyzes triglycerides into glycerol and fatty acid components until only glycerol is left, which occurs through three enzymatic reactions [6]. The basic enzymes for lipolysis are adipose triglyceride lipase (ATGL), hormone-sensitive lipase (HSL), and monoacylglycerol lipase (MGL). ATGL hydrolyzes triacylglycerol (TAG) to diacylglycerol (DAG) and one fatty acid (FA). HSL converts DAG to monoacylglycerol (MAG) and one FA, and MGL hydrolyses MAG to produce glycerol and the third FA. MGL is a recently discovered cancer-related protein in addition to its role as a lipid-degrading enzyme factor. Studies have demonstrated that overexpression of MGL inhibits cancer cell growth [7].

A Food and Drug Administration (FDA)-approved product containing semaglutide, a glucagon-like peptide 1 (GLP-1) receptor agonist that targets brain regions, has side effects including vomiting, nausea, constipation, diarrhea, dizziness, insomnia, and liver damage [8, 9]. These drugs induce the pituitary gland to suppress appetite and are not directly related to the breakdown of fat cells or lipolysis. Despite advances in chemical therapy, obesity is still difficult to treat due to a lack of understanding of the pathological and physiological mechanisms of lipolysis. Ultrasound technology has been developed as diagnostic and therapeutic agents in wide range of diseases including immunotherapy for cancer, cataract, drug delivery, and tissue regeneration [10-15]. Ultrasound may be used to treat obesity by directly breaking down fat cells and / or stimulate brain regions responsible to control appetite and food consumption.

Low intensity pulsed ultrasound (LIPUS) stimulation uses non-invasive, non-ionized mechanical energy transfer to stimulate biological objects from cultured cells to whole organs in living animals and humans. Therefore, LIPUS stimulation has emerged as a valuable biomedical tool in several preclinical studies including neuromodulation, pain relief, inflammatory mediation and tissue and bone healing with potential applications in clinics. Ultrasonic stimulation of various types of cells in culture is actively studied in many groups because it is known that mechanical force plays a key role in regulating various cellular functions such as stem cell differentiation and proliferation [16], cancer cell death [17], and the regulation of cell functions [18]. In clinical trials, high intensity focused ultrasound with the intensity of over 5 W/cm^2^ directly ablates the region of interests to induce immune stimulatory effects [19]. LIPUS may induce inflammatory responses for cancer immunotherapy, which may be introduced as the next generation approach to reduce immune suppression and drug resistance [17, 20]. Although LIPUS stimulation has been widely used in research laboratories and clinics, the fundamental mechanism of how fat cells respond to LIPUS stimulation has not been studied at molecular level.

In this study, we investigated the function of the LIPUS stimulation on biophysical metabolism of cultured adipocytes. Although chemical therapies are frequently used to reduce obesity in clinics, they sometimes induce side effects. Our goal is to explore potential opportunities as the next generation therapy for obesity using LIPUS stimulation with minimum side effects after understanding the fundamental mechanism of lipolysis of adipocytes on LIPUS stimulation. Adipocytes were exposed to 2 MHz LIPUS for 10 minutes every day for three days. Three different intensities of LIPUS were applied to cultured adipocytes to compare the changes of lipolytic factors after LIPUS stimulation at RNA and protein levels by quantitative reverse transcription polymerase chain reaction (RT-qPCR) and Western blot. We explored genes affected by LIPUS stimulation using transcriptomic profiling and pathway analysis by RNA-sequencing (RNA-Seq) analysis. We selected target genes with high fold changes and confirmed their expression in adipocytes *in vitro* using immunofluorescence staining. Based on our studies, we will ultimately aim to utilize LIPUS stimulation for *in vivo* and potentially clinical applications after further understanding of the mechanisms underlying the regulation of adipocyte degradation.

## 2. Materials and methods

### 2.1. Cell culture and differentiation

3T3-L1 (American Type Culture Collection (ATCC, USA)) cells were cultured in DMEM supplemented with 10% fetal bovine serum (FBS, Thermo Fisher Scientific, USA) and 1% penicillin/streptomycin (P/S, Thermo Fisher Scientific, USA) at 37 °C in a humidified atmosphere with 5% CO_2_. For adipogenic differentiation, 3T3-L1 cells were exposed to differentiation medium. Differentiation medium is DMEM supplemented with 10% FBS, 1% P/S, 2.5 mM isobutyl methylxanthine (Sigma, USA), 1 μM dexamethasone (Sigma, USA), and 1 μg/ml insulin (Sigma, USA) for 3 days and then maintained in medium containing 1 μg/ml insulin for 4 days. Then, it was changed to DMEM medium and used for the experiment.

### 2.2. LIPUS stimulation system and protocol

A 2 MHz single element transducer for LIPUS stimulation was developed using piezoelectric materials (PZT-4, Boston piezo-optics, USA) by following a protocol in our lab [21-23]. The aperture and the focus of the PZT-4 were 40 mm in diameter and 50 mm, respectively. The *f-number* of the PZT-4 was 1.25. PZT-4 was place at the distal end of a machined brass housing using epoxy. Hot wire was linked to a SMA connector and chrome / gold (Cr / Au) with the thickness of 50 nm and 400 nm were sputtered on the front side of the aperture of PZT-4 placed on the bass housing for ground connection.

For LIPUS stimulation, the developed transducer was operated in a pulsed mode with a 100-cycle sine wave burst with peak-to-peak voltages (V_pp_) of 31.6 V_pp_ and 56.9 V_pp_ and 1 ms pulse repetition time (PRT) (Fig. 1a, c). The sine wave burst was generated by a function generator (AFG31252, Tektronix, USA) and amplified by a 50 dB power amplifier (525LA, E&I ltd., USA) (Fig. 1a). The whole system was fixed on an optical board. A water cuvette was used to couple the transducer and a cell culture dish. Three-axis computer-controlled position system (ILS150CC, Newport Corp., USA) was used to scan 30 mm by 30 mm area to stimulate 3T3-L1 cells in a 35 mm dish (Fig. 1a, b) [24, 25]. A custom LabView (National Instruments, USA) computer program was used to control the positioning system to repeatedly scan the same area seven times for 10 minutes. LIPUS stimulation was repeated for three days.

**Fig. 1.**
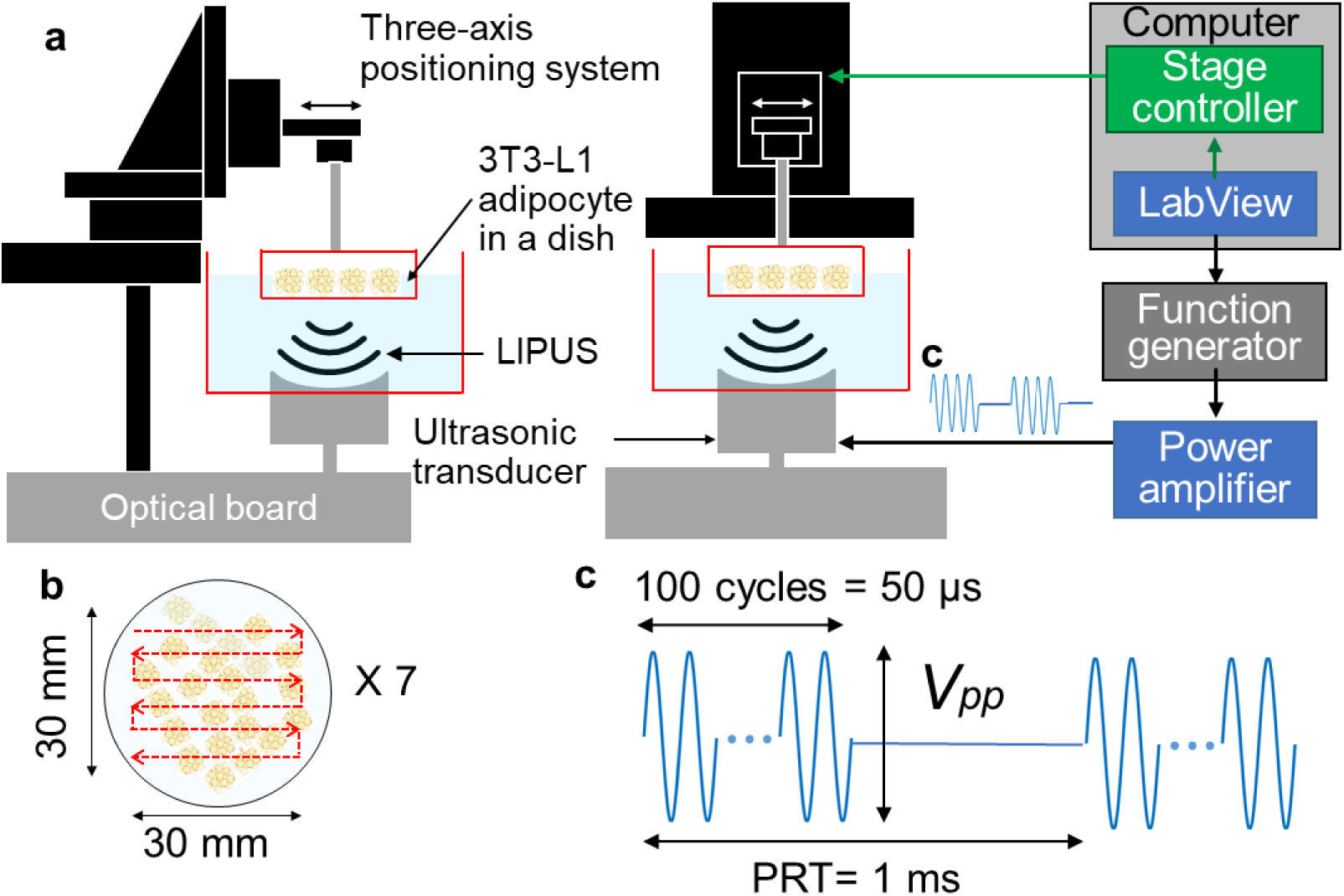
Schematic diagram showing the ultrasound system. **a**. Three-axis computer-controlled position system for low intensity pulsed ultrasound (LIPUS) stimulation was controlled by a LabView computer program (**b)** to scan 30 mm by 30 mm area of a dish, where 3T3-L1 adipocytes were cultured. **b**. Scan was repeated seven time and it took approximately 10 minutes. **c**. The 2 MHz ultrasonic transducer was triggered by pulsed electrical signal with peak-to-peak voltage (V_pp_) of 31.6 V_pp_ and 56.9 V_pp_, 100-cycle sine waves, and pulse repetition time (PRT) of 1 ms.

The acoustic output such as acoustic pressure and spatial peak temporal average intensity (*I*_*SPTA*_) of the 2 MHz ultrasonic transducer was measured with a needle hydrophone (HGL-0085, Onda) using the same three-axis positioning system [25, 26].

### 2.3. RT-qPCR

Total RNA from sample was isolated using TRIzol reagent (Thermo Fisher Scientific, USA) according to the manufacturer’s instructions. Converted to cDNA using a high-capacity cDNA reverse transcription kit (Applied Biosystems, USA). Quantitative PCR was performed for 45 cycles using SYBR Green Supermix (BioRad, USA) and CFX Real-time PCR System (BioRad, USA), and the fold change of all samples was calculated using the Ct method. cDNA was amplified using the primers listed in Table S1.

### 2.4. Western blot

For Western blot analyses, cells were lysed in RIPA Lysis and Extraction Buffer (Thermo Fisher Scientific, USA) containing SIGMAFAST Protease Inhibitor Cocktail (Sigma, USA) and PhoSTOP phosphatase inhibitors (RocheProtein concentration was quantified using Pierce BCA Protein Assay Kit (Thermo Fisher Scientific, USA). Protein samples were separated on SDS– PAGE gel and transferred to PVDF membrane (BioRad, USA). Membranes were incubated in blocking buffer (Tris-buffered salinewith 0.1% Tween 20 (TBS-T) containing 5% non-fat dry milk) and then sequentially incubated with primary antibody at 4 °C overnight and horseradish peroxidase-conjugated secondary antibody at room temperature for 1 h. Antibodies were diluted in the blocking buffer. The primary antibodies used for Western blot analysis are listed in Table S2. Western blot images were captured with the chemiluminescence imaging system (BioRad, USA). Quantification of western blot was performed using the National Institutes of Health (NIH) ImageJ software. Unedited western blot membrane images are available as source data.

### 2.5. RNA-seq analysis

RNA samples were extracted using the TRIzol reagent following the manufacturer’s protocol. Samples with RIN >10 were used for RNA-seq library preparation at the Notre Dame Genomics Core Facility. RNA-seq libraries were prepared with NEBNext Ultra II Directional RNA Library Prep Kit (mRNA) and NextSeq Illumina Sequencing. Raw RNA-seq data were deposited with Bioinformatics consulting (University of Notre Dame). Raw sequences were trimmed of adapters with Trimmomatic version 0.39 [27] and assessed for quality with FastQC v 0.11.8 [28]. Trimmed sequences were aligned to the mouse genome, Ensembl build Mus_musculus. GRCm39 using Mus_musculus.GRCm39.104 version annotations and HISAT2 version 2.1.0 [29]. Corresponding alignments were sorted with SAMtools version 1.9 [30]. Read counts were generated with HTSeq-count version 0.11.2[31] and were merged with a python script [32]. Subsequent statistics were completed in R (R Core Team, 2014) implementing the EdgeR library [33-36].

### 2.6. Immunofluorescence (IF)

To stain the lipid droplets of 3T3-L1 cells, BODIPY (Thermo Fisher Scientific, USA) was stained for 15 minutes, washed 3 times with PBS, and then the cells were fixed with 4% Paraformaldehyde (PFA) (Sigma, USA) at room temperature. cell was then blocked with 5% BSA at room temperature for 1 hour and incubated with mitochondrially encoded cytochrome c oxidase III (Mt-co3) S1P1 and macrophage receptor with collagenous structure (MARCO) primary antibody. The next day, cell was washed 3 times with PBS and incubated with the following secondary antibodies. After 45 min incubation at room temperature, PBS washed and then mounted with DAPI (4′,6-diamidino-2-phenylindole, Sigma, USA), which stains the nuclei of immune cells to visualize the nuclei. Slides were imaged with a Nikon microscope (Nikon Eclipse Ti2, Japan) and Nikon confocal microscope (A1R-MP Laser Scanning Confocal Microscope, Japan).

### 2.7. Triglyceride (TG) assay

Lysis cells were analyzed for triglyceride content using a commercially available colorimetric assay from Triglyceride Colorimetric Assay Kit (Cayman, USA). 10 ul of each sample was placed in each well, then 150 ul of enzyme buffer was added and the absorbance was read at 540 nm. A standard curve was run on each assay plate using recombinant proteins in serial dilutions.

### 2.8. Glycerol assay

Glycerol was quantified using the Sigma-Aldrich Glycerol Assay Kit (Sigma, USA) colorimetric assay according to the supplier’s instructions. Samples were diluted with sterile water to a detectable concentration within the standard curve.

### 2.9. Statistical analysis

Statistical analyses were performed using GraphPad Prism 9 software (GraphPad Software, La Jolla, CA, USA). Heatmaps were generated by Origen program. Statistical significance was established at a value of *p* < 0.05 and *p* < 0.01.

## 3. Results and discussion

### 3.1. LIPUS stimulation induces lipolysis in adipocytes differentiated 3T3-L1 cells

Acoustic pressure and *I*_*SPTA*_ at the focus of the 2 MHz ultrasonic transducer for LIPUS stimulation were measured to be 0.31 MPa and 370 mW/cm^2^ when V_pp_ was 31.6 V_pp_ and 0.55 MPa and 444 mW/cm^2^ when V_pp_ was 56.9 V_pp_. The mechanical indexes of V_pp_ of 31.6 V_pp_ and 56.9 V_pp_ were estimated to be 0.22 and 0.39. These values are below the FDA safety limit for diagnostic ultrasound [37].

3T3-L1 cells are white adipocytes and the most widely used cell line in biological studies of obesity and adipose tissue. We hypothesized that ultrasound stimulation may lead lipolysis. We searched for the ranges of ultrasound intensities to safely simulate adipocytes to induce lipolysis from previous studies [38]. Lipolysis is mainly catalyzed by three major lipases: ATGL, HSL, and MGL, which belong to the standard pathway of adipocyte lipolysis [39]. The expression levels of several lipolytic genes were confirmed in adipocytes by RT-qPCR (Fig. 2a). Gene expression levels indicated that LIPUS stimulation with 370 mW/cm^2^ and 444 mW/cm^2^ upregulated the expression of lipolysis-related genes. Perilipin 5 (Plin5) was significantly increased at 370 mW/cm^2^ and 444 mW/cm^2^ LIPUS stimulations compared to the control group. Plin5 is a lipid droplet-related protein that is thought to control intracellular lipid metabolism in highly oxidized tissues and in response to phosphorylation by protein kinase A (PKA) [40]. Perilipin1 (Plin1) was significantly increased at 444 mW/cm^2^ LIPUS stimulation compared to the control group. The increase of cyclin D1 (Ccnd1) was not significant.

**Fig. 2.**
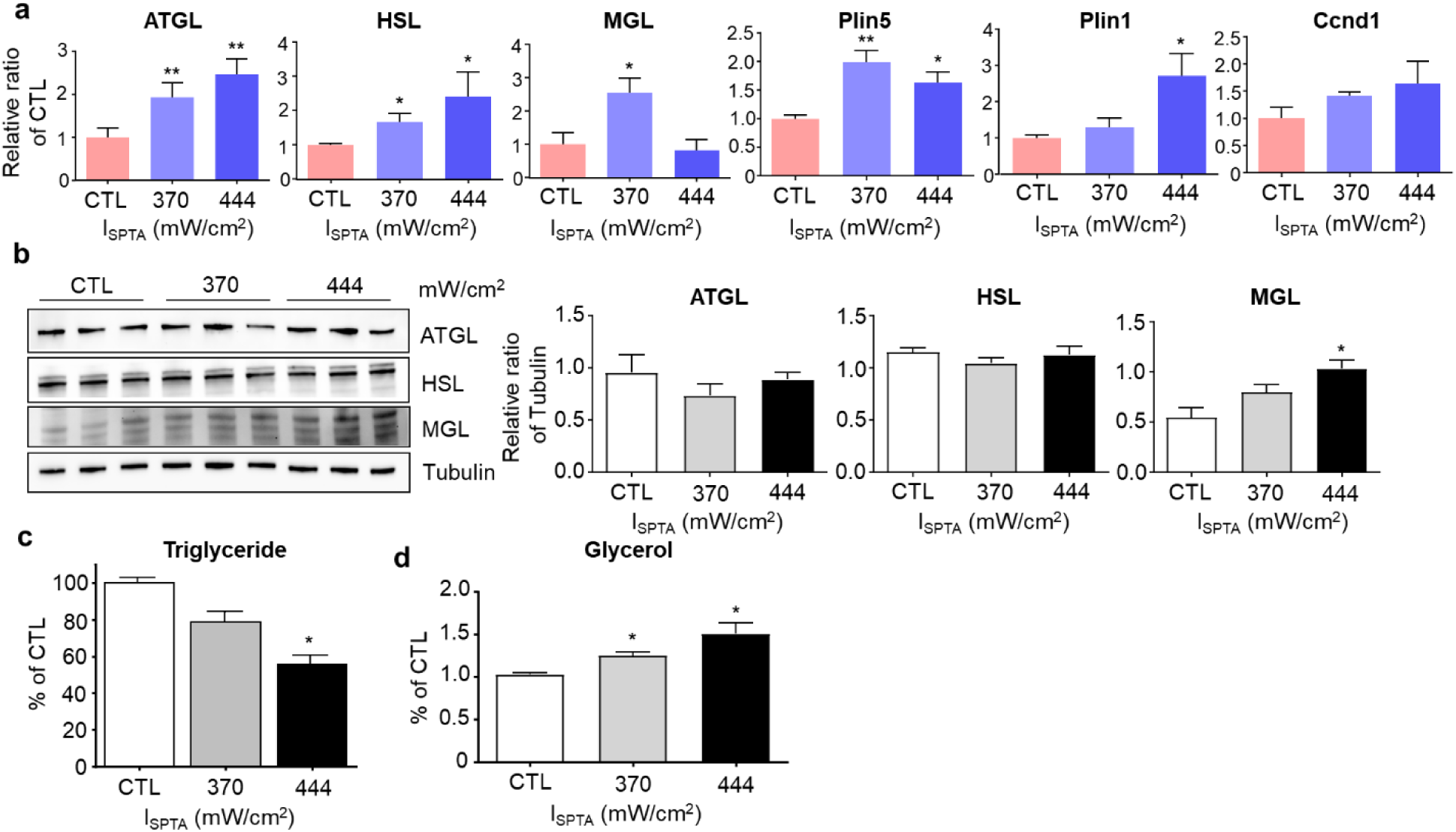
Effect of lipolysis in adipocytes differentiated from 3T3-L1 cells by LIPUS stimulation. **a**. RT-qPCR analysis shows the changes of the level of expressions of adipose triglyceride lipase (ATGL), Hormone-sensitive lipase (HSL), monoacylglycerol lipase (MGL), Perilipin5 (Plin5), Perilipin 1 (Plin1), and Cyclin D1 (Ccnd1). Spatial peak temporal average intensity (*I*_*SPTA*_) of 370 mW/cm^2^ and 444 mW/cm^2^ were applied and compared with control group (CTL) with no LIPUS stimulation. **b**. Western blot analysis shows that only MGL protein is significantly upregulated by LIPUS stimulation. **c**. The triglyceride level of 3T3-L1 cells is significantly decreased under LIPUS stimulation with 444 mW/cm^2^, measured by triglyceride kit. **d**. The level of glycerol is increased under LIPUS stimulation with 370 mW/cm^2^ and 444 mW/cm^2^. Glycerol is a substance hydrolyzed from triglycerides, suggesting that fats are broken down to glycerol. All experiments were performed three times (n= 3). Error bars indicate ± standard error of the mean (SEM). We used paired t-test to calculate the p values and *p < 0.05, **p < 0.01 were considered statistically significant.

To determine the effect of LIPUS stimulation on protein expression, we performed Western blot analysis of proteins in the groups with and without LIPUS stimulation. Western blot analysis indicated that LIPUS stimulation increased the levels of MGL (Fig. 2b). However, we observed insignificant changes of ATGL and HSL between control and LIPUS stimulation groups (Fig. 2b). Based on this observation, we may use LIPUS stimulation as a potential and alternative therapy for obesity with lower side effects due to different stimulation mechanism of lipolysis compared to chemical stimulations. Chemical stimulations increase the activation of all three layers of ATGL, HSL, and MGL at both RNA and protein levels during lipolysis, which may cause side effects [8]. In chemical stimulations, treatment with the peroxisome proliferator-activated receptor (PPAR) agonist, rosiglitazone, simultaneously increased ATGL, HSL, and MGL in white adipose tissue (WAT) at RNA level [41]. Lipolysis activators include catecholamines such as epinephrine and isoproterenol, adrenocorticotropic hormone, and glucagon. All of these substances increase cAMP levels, which indirectly activate ATGL, HSL, and MGL [42]. Adrenergic stimulation of isoproterenol slightly altered ATGL and HSL proteins in primary rat adipocytes. It also significantly promote glycerol release [43]. It was reported that HSL was increased at gene and protein levels by the treatment of epigallocatechin gallate (EGCG), which may be obtained from green tea [44].

We found 21% and 44% reduction of triglyceride in stimulated cells with 370 mW/cm^2^ and 444 mW/cm^2^ LIPUS, compared to the control group. (Fig. 2c). The decreased triglyceride levels observed in cells treated with LIPUS may be due to a combination of decreased basal adipogenic activity and increased basal lipolytic activity in addition to delayed terminal adipocyte differentiation. 370 and 444 mW/cm^2^ LIPUS stimulation increased the glycerol content in adipocytes by 5%, compared to the control group. Collectively, these results indicate that LIPUS stimulation may contribute the activation of lipolysis.

### 3.2. Gene profiling after LIPUS stimulation

After confirming gene regulation by LIPUS stimulation, RNA-sequencing (RNA-seq) was performed to explore the effects of LIPUS on broad range of genes. We performed RNA-seq profiling of control group and experimental groups stimulated by LIPUS with 370 mW/cm^2^ and 444 mW/cm^2^ intensities. Based on the 14,572 gene probes surveyed in RNA-seq, we generated volcano plots and MA plots comparing the effect of different LIPUS stimulations and found evidence that LIPUS stimulation has impact on gene expression in lipolysis and autophagy (Fig. 3f and g). We generated a heat map (Fig. 3a) and completed an ingenuity pathway analysis which identified 224 canonical pathways affected by LIPUS stimulation. 7,206 genes were upregulated, and 7,367 genes were downregulated at 370 mW/cm^2^ of LIPUS stimulation (Fig. 3a). 7,367 genes were upregulated in the 444 mW/cm^2^ LIPUS stimulation group, while 7,205 genes were downregulated (Fig. 3a). A total of 160 genes were differentially regulated in adipocytes between 370 mW/cm^2^ and control and 444 mW/cm^2^ and control. For example, cells stimulated by 370 mW/cm^2^ LIPUS were highly enriched fatty acid oxidation III signaling (Fig. 3d). In comparison, cells stimulated by 444 mW/cm^2^ LIPUS were enriched in aryl hydrocarbon receptor (AHR) signaling (Fig. 3e). Fatty acid oxidation III is esterified with complex lipids such as TAG or via carnitine palmitoyl transferase (CPT). Upregulation of fatty acid oxidation III is related to lipid oxidation and AHR regulates pathological conditions ranging from tissue homeostasis and inflammatory diseases to neoplastic diseases, which may promote lipolysis [45]. We chose Mt-co3 gene, which is one of the genes in fatty acid oxidation III, S1P1 signaling gene, and MARCO, which is one of the genes involved in inflammasome pathway for further investigation based on high fold changes in volcano and MA plots (Fig. 3d - g).

**Fig. 3.**
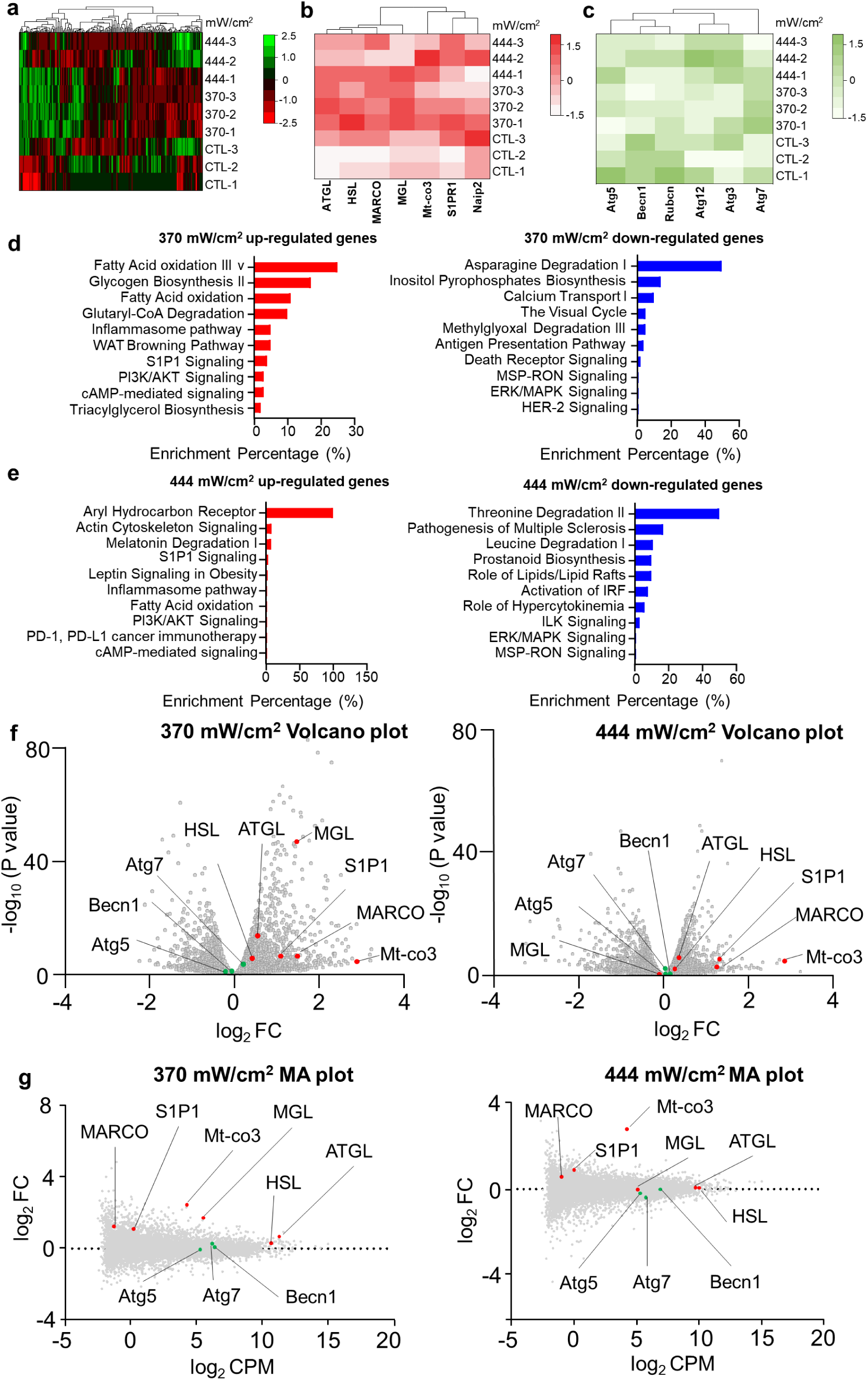
RNA-seq analysis of genes and signaling pathways after LIPUS stimulation. **a**. Heat map shows differentially regulated genes after LIPUS stimulation with 370 mW/cm^2^ and 444 mW/cm^2^. Gene expression is shown in normalized log_2_ counts. **b**. Heat map shows lipolysis-related genes. Red and white colors indicate the expression level of differentially expressed genes. **c**. Heat map shows autophagy-related genes. Dark green and white colors indicate the expression level of differentially expressed genes. **d** and **e**. Gene pathway analysis were performed for differentially expressed genes. Top 10 up- and down-regulated pathways are shown for LIPUS stimulation with 370 mW/cm^2^ and 444 mW/cm^2^. **f**. Volcano plots show the fold change and p-value of individual genes after LIPUS stimulation with 370 mW/cm^2^ and 444 mW/cm^2^. The log_2_ fold change (FC) ratios of 370 mW/cm^2^ vs. control and 444 mW/cm^2^ vs. control were plotted with respect to -log_10_(p-value). **g**. MA plots show the relationship between average concentration (log_2_ CPM) and fold-change (log_2_ FC) across the genes. Each gene is represented by a gray dot. Genes related to lipolysis from **b** are indicated as red dots, and genes related to autophagy from **c** are shown in green dots.

### 3.3. LIPUS stimulation induces mitochondria activation and immune regulation in adipocytes

We investigated gene expression and protein expression of Mt-co3, S1P1, and MARCO based on RNA-seq results. At RNA level, Mt-co3, S1P1, and MARCO increased under 370 mW/cm^2^ LIPUS (Fig. 4a). LIPUS stimulation activates mitochondria and immune related genes. Western blot analysis indicates that Mt-co3, S1P1 and MARCO were increased as LIPUS intensity increases (Fig. 4b).

**Fig. 4.**
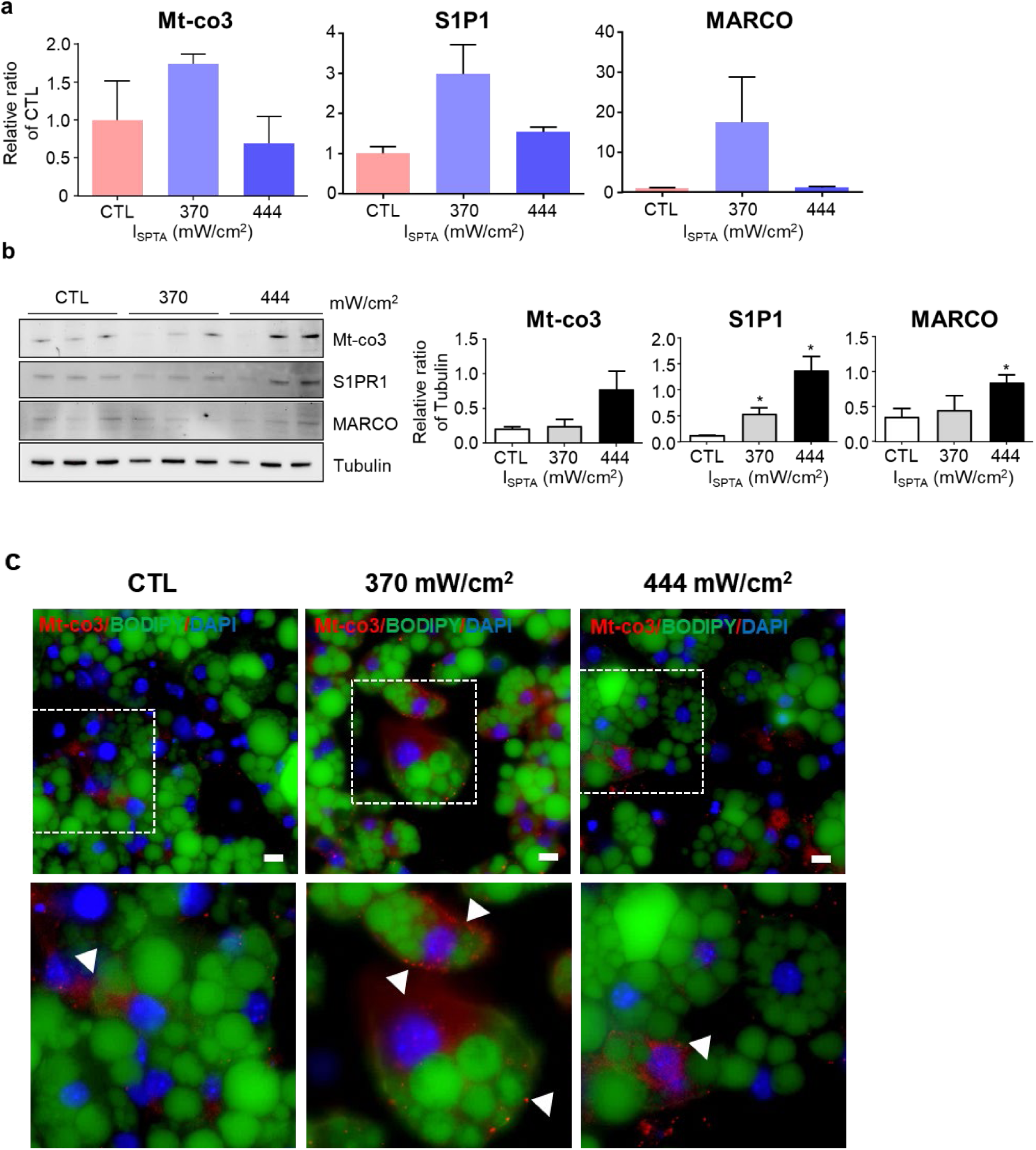
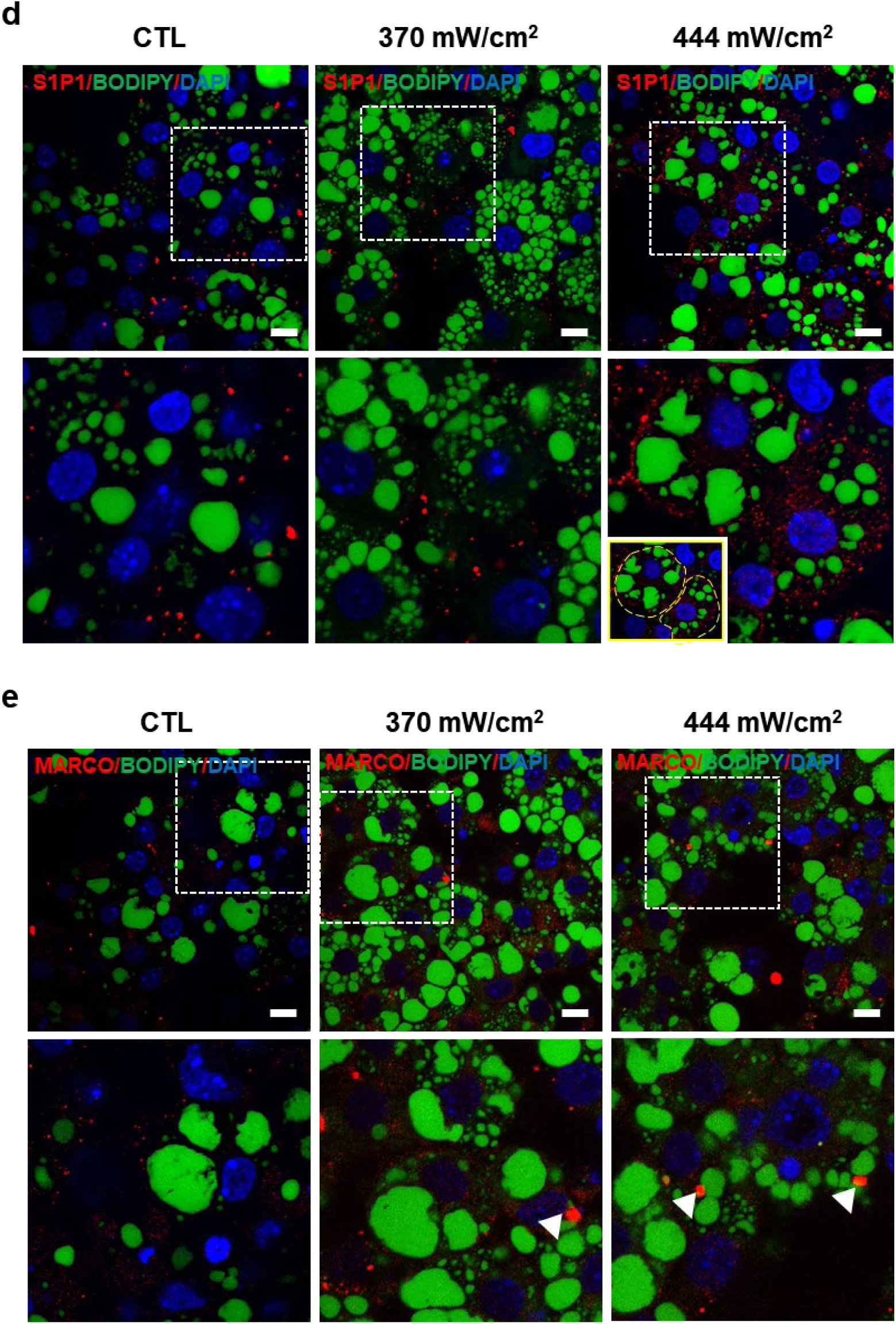
LIPUS stimulation induces mitochondria activation and immune regulation in adipocytes. **a**. mRNA levels were quantified by RT-qPCR. The mRNA level of individual mitochondrially encoded cytochrome c oxidase III (Mt-co3), sphinosine-1-phosphate (S1P1), and macrophage receptor with collagenous structure (MARCO) genes was normalized to the mRNA level of the corresponding gene in control cells, which was set to 1. The changes are not statistically significant. **b**. Western blot analysis of Mt-co3, S1P1 and MARCO expression in adipocytes differentiated from 3T3-L1 cells treated with 370 mW/cm^2^ and 444 mW/cm^2^ LIPUS stimulation shows significant upregulation of S1P1 and MARCO while Mt-co3 shows small changes. Error bars indicate ± one standard deviation. We used paired t-test to calculate the p values and *p < 0.05 was considered statistically significant (n = 3). Immunofluorescence staining of **c**. Mt-co3, **d**. S1P1 and **e**. MARCO with 370 mW/cm^2^ and 444 mW/cm^2^ LIPUS stimulation. Mt-co3 expressions in the mitochondria are indicated by arrow heads in **c**. S1P1 in **d** is coupled with G protein-coupled receptors located on the cell plasma membrane. Yellow dashed lines in inset in **d**. indicate S1P1 protein on cell boundary. MARCO is related to macrophage phagocytosis and the expression is indicated by arrow heads in **e**. Green and blue stainings represent lipid drop and nucleus, respectively. Scale bar:10 µm

Immunostaining images of Mt-co3, S1P1, and MARCO confirmed that LIPUS stimulation promotes the activity of mitochondria in adipocytes, S1P1, and MARCO. Increased expression of Mt-co3, S1P1, and MARCO were observed indicated as white arrow heads and red fluorescence in Figures 4c-e. S1P1 expression with red fluorescence indicates the localization of S1P1 on cell plasma membrane (yellow dashed line in inset in Fig. 4d). Mitochondria play an important role in regulating adipocyte fatty acid storage by regulating fatty acid breakdown through oxidation as well as synthesis. S1P1 is a lipid mediator that can activate G protein-coupled receptors (GPCR) [46]. Macrophage-mediated phagocytosis plays an important role in maintaining tissue homeostasis and is the first line of defense against invading pathogens. The phagocytic capacity of macrophages decreases when MARCO is knocked down [47]. Based on our results, LIPUS stimulation may enhance the expression of mitochondrial genes and activity, the activation of GPCR, and the digestion of lipid droplets into small pieces. BODIPY-green and DAPI were used to stain lipid droplets and nucleus of adipocytes differentiated from 3T3-L1 cells, respectively (Fig. 4c-e).

### 3.4. Effect of LIPUS stimulation on autophagy response of adipocytes

LIPUS stimulation upregulated MGL protein while it did not upregulate the expression of ATGL and HSL proteins (Fig. 2b). Despite this observation, lipid content of adipocytes decreased (Fig. 2 c and d). Therefore, we hypothesized that LIPUS stimulation may activate an alternative pathway for lipid consumption. Lipolysis was considered an enzymatic activity that liberates fatty acids as metabolic fuel. Recent studies have shown that novel substrates containing various lipid compounds, such as fatty acids and their derivatives, release lipolysis products that act as signaling molecules and transcriptional regulators [48]. Autophagy is a pathway that mediates vacuolar degradation and protein recycling. It has also been reported as a novel regulator of lipolysis, which plays an important role in cell physiology and acts as a lysosomal lipolysis pathway [49]. We investigated the upregulation and downregulation of some of autophagy related genes (ATG) under LIPUS stimulation.

To measure autophagy markers, ATGs were measured by RT-qPCR and Western blot analysis. Beclin1(Becn1) showed insignificant changes after LIPUS stimulation at RNA and protein levels (Fig. 5a and b). While Atg5 expression at transcriptional level was not statistically significant, its expression at translational level was statistically significant under LIPUS stimulation with 370 mW/cm^2^ and 444 mW/cm^2^ (Fig. 5a and b). Based on the results, we observed changes of ATGs at RNA and protein levels, which may be an indication of LIPUS mediated regulation of autophagy related genes of white adipocyte. Although there is a growing evidence for roles of autophagy in lipid storage and degradation, further studies are needed to fully understand the mechanisms of autophagy under LIPUS and investigate the potential therapeutic applications by targeting autophagy in wide ranges of diseases.

**Fig. 5.**
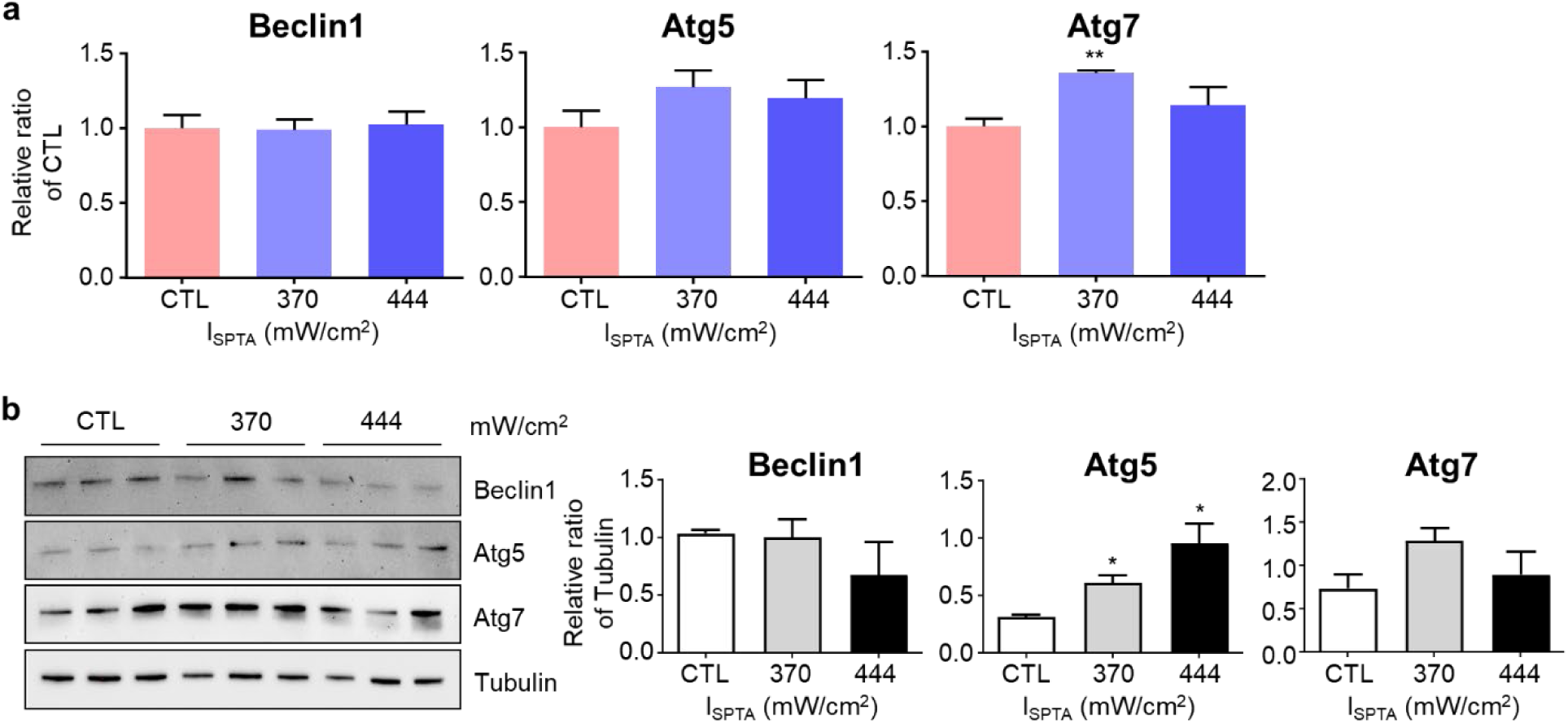
Effect of LIPUS stimulation on autophagy related genes of adipocytes. **a**. mRNA levels were quantified by RT-qPCR. The mRNA level of individual autophagy related gens (ATG) genes was normalized to the mRNA level of the corresponding gene in control cells, which was set to 1. Atg7 shows significant changes while Beclin1(Becn1) and Atg5 are not significantly changed under LIPUS stimulation. **b**. Western blot analysis of autophagy marker expressions shows significant changes in Atg5 proteins. Error bars indicate ± SEM. We used paired t-test to calculate the p values and *p < 0.05 and **p < 0.01 were considered statistically significant. (n = 3).

## 4. Conclusions

The mechanisms involved in various biological effects on adipocytes by LIPUS stimulation have not been largely studied. This study investigated the effects of LIPUS stimulation on lipolysis of adipocytes differentiated from 3T3-L1 cells at molecular level by applying different levels of acoustic intensities. LIPUS stimulation particularly upregulates MGL proteins among three lipolytic factors. Compared to chemical stimulation, LIPUS stimulation induces distinct activation of lipolysis related proteins which may potentially be used as a next generation therapy to reduce fat stably and with minimal side effects. Besides, we observed the regulation of various genes that promotes mitochondria activation, immune regulation, and cell division by LIPUS with different acoustic intensities. In addition, LIPUS stimulation induces the upregulation of autophagy related genes, which may be a potential evidence of autophagy mediated clearing of substances produced by lipolysis to maintain the homeostasis of adipocytes.

## Acknowledgments

We thank Melissa Stephens and Joseph Sarro for RNA-sequence analysis at Genomics and Bioinformatics Core Facility (University of Notre Dame). This work was supported in part by the National Institutes of Health (NIH) under Grant No. GM120493, the National Science Foundation (NSF) under Grant No. CBET 1943852, and the Harper Cancer Research Institute, CCV grant.

## Author contributions

S. Kim and S. Yoon conceived the idea. S. Kim and S. Yoon designed, performed, and analyzed experiments. S. Kim and S. Yoon wrote and edited the manuscript. S. Yoon supervised the whole procedure.

